# Genomic signatures of selection are enriched in differentially expressed genes in sticklebacks adapting to contrasting environments

**DOI:** 10.64898/2026.02.22.707242

**Authors:** Alexander Kwakye, Natalie Dzikowski, Matthew A. Wund, Krishna R. Veeramah

## Abstract

Whole genome scans have identified numerous adaptive alleles in many species; however, linking these alleles to specific phenotypes remains a major challenge. A promising alternative to direct genotype–phenotype mapping, particularly given the complexities introduced by epistasis, pleiotropy, and environmental variability, is to assess whether differentially expressed genes are enriched in regions of genetic divergence between populations adapted to contrasting environments. Here, we study gene expression patterns in threespine stickleback populations adapting to contrasting environments (marine vs freshwater) and investigate how signatures of selection interact with patterns of gene expression. We performed transcriptomic experiments of the brain and gill tissues of wild-caught sticklebacks sampled from one marine and two freshwater environments using TagSeq. We found that differentially expressed genes in the freshwater environments are enriched for single nucleotide polymorphisms (SNPs) previously identified to be involved in rapid adaptation and F*_ST_* outliers. A majority of these SNPs were located in *cis*-regulatory regions with predicted low to moderate effects on protein function and structure, although we found a high-impact SNP in the gene *col8a1b*. Genes such as *pvalb4* and *acsl4a*, involved in calcium regulation in the gill and fatty acid metabolism in the brain, respectively, were enriched with SNPs showing signatures of selection. By linking signatures of selection to tissue-specific gene expression patterns, our study bridges the gap between genomic divergence and the molecular mechanisms underlying physiological adaptation to new environments.

**Significance Statement:** Understanding how genetic variation translates into adaptive traits remains a central challenge in evolutionary biology. While whole-genome scans routinely identify candidate adaptive alleles, connecting these variants to functional phenotypes is complicated by epistasis, pleiotropy, and environmental effects. Here, we integrate signatures of selection with tissue-specific gene expression in threespine stickleback adapting to contrasting environments (marine and freshwater). We demonstrate that differentially expressed genes in freshwater populations are enriched for previously identified adaptive SNPs and F*_ST_* outliers, many of which are located in *cis*-regulatory regions with predicted low to moderate functional effects. Notably, we identify a high-impact variant leading to a premature stop codon in *col8a1b* and highlight genes such as *pvalb4* and *acsl4a* that link selection to key physiological processes, including ion regulation in gills and fatty acid metabolism in the brain. By connecting genomic divergence to regulatory and tissue-specific expression changes, this work provides a mechanistic framework for understanding how natural selection shapes complex physiological adaptation.

## Introduction

In the last 25 years the emergence of genomic-scale data has enabled the discovery of numerous loci across many species that demonstrate signals consistent with positive selection(Radwan & Babik 2012; Campagna & Toews 2022; Hao et al. 2024; Yang et al. 2024; Peng et al. 2024; Jones et al. 2012; Roberts Kingman et al. 2021; Moreira & Smith 2023; Massey & Wittkopp 2016; Bomblies & Peichel 2022; Kerem et al. 1989; Linnen et al. 2009; Colosimo et al. 2004; Lamichhaney et al. 2015). While some of these loci have been shown to also have large phenotypic effects(Linnen et al. 2009; Colosimo et al. 2004; Lamichhaney et al. 2016, 2015), in most cases the traits influenced by adaptive alleles have remained unidentified. Theoretical models have long suggested that adaptation is often driven by many alleles of small effects(Fisher 1958, 1999; Kimura 2012). More recent empirical work, particularly QTL-based studies, have frequently identified loci of moderate to large effect(Colosimo et al. 2005; Schluter et al. 2021; Fang et al. 2020; Hoekstra et al. 2006), though such loci may be disproportionately detectable relative to small-effect variants. Consequently, a major challenge moving forward is to link these putative adaptive alleles of small effect to the phenotypes they influence.

Mapping genotypes to specific phenotypes, especially alleles of small effect, remains daunting for several reasons. First, environmental variables can substantially influence gene expression, altering the relationship between phenotype and the underlying genetic variation. Furthermore, the genetic background can modulate an allele’s effect through mechanisms such as epistasis(Stern et al. 2022) or pleiotropy(Rennison & Peichel 2022; McKay et al. 2003). Another layer of complexity lies in how phenotypes are defined(Ahnert 2017); phenotypes may encompass amino acids or the protein residues they encode, as well as broader molecular pathways, physiological processes, or morphological traits that arise from these molecular entities or processes. The more hierarchical levels separating the genotype and associated phenotype of interest, the more complex the mapping relationship is likely to be. In addition, large sample sizes are required to detect alleles of small effects when conducting QTL studies(Mackay et al. 2009).

A promising approach that potentially side-steps at least some of these complexities involves the estimation of the enrichment of differentially expressed genes in regions of genetic divergence between populations inhabiting contrasting environments(Mack et al. 2023; Rodríguez-Ramírez & Peichel 2025; Verta & Jones 2019; Forsberg et al. 2025). This approach is based on an assumption that genetic signatures of adaptation found in or nearby differentially expressed genes are involved in gene expression divergence between populations in contrasting environments. By design, this approach targets variants found in gene bodies and *cis*-regulatory regions, rather than *trans*-regulatory variants or even more distantly-acting variants that may be in linkage disequilibrium with target variants.

Threespine stickleback (*Gasteroteus aculeatus*) is an ancestral marine(Bell et al. 2009) fish that is found along the coastal regions of the Northern hemisphere. There are many ecotypes of this fish, which are often considered to reside in pairs of contrasting environments, such as marine – freshwater, benthic–limnetic, and lake– stream habitats(Reid et al. 2021). The marine ecotype has repeatedly adapted to new freshwater environments for at least the past 10 million years(Bell & Foster 1994). Adaptation to freshwater environments is characterized by rapid phenotypic changes, which are observed within a few generations(Heuts 1947; Bell & Richkind 1981; Bell et al. 2004), as well as rapid genomic divergence(Colosimo et al. 2004; Cresko et al. 2004; Marchinko et al. 2014; Jones et al. 2012; Roberts Kingman et al. 2021; Kwakye et al. 2025). Changes in more than 300 loci likely underlie freshwater adaptation as demonstrated by rapid shifts in their respective allele frequencies when marine fish are introduced into freshwater environments(Jones et al. 2012; Hohenlohe et al. 2012; Roberts Kingman et al. 2021). However, with the exception of a handful of loci, their functional role in freshwater adaptation is typically unknown(Reid et al. 2021).

It is hypothesised that *cis*-regulatory changes that alter gene expression likely underlie the genetic mechanism by which many of these loci drive freshwater adaptation(Jones et al. 2012; Mack et al. 2023). Using F1 marine-freshwater hybrids of threespine stickleback, Mack et al. 2023(Mack et al. 2023) provided evidence that allele-specific expressed genes are enriched in regions of marine-freshwater divergence(Mack et al. 2023). Differentially spliced genes between marine and freshwater ecotypes of threespine stickleback have also been found to be enriched in these genomic regions of marine-freshwater divergence(Rodríguez-Ramírez & Peichel 2025). However, it remains unclear if these gene expression changes reflect those observed in nature. Most studies that have quantified gene expression changes have used lab raised threespine stickleback(Mack et al. 2023; Rodríguez-Ramírez & Peichel 2025; Verta & Jones 2019). Laboratory conditions control biological noise but eliminate the influence of environmental variability, which is a crucial factor that affects genotype–phenotype relationships. Environmental volatility can also be impactful in the molecular processes that allow an organism to adapt to its local habitat.

The gill and brain play important roles in physiological and behavioral phenotypes involved in adaptation to new freshwater environments by marine threespine stickleback(Di Poi et al. 2016). The gill is a key osmoregulatory organ important for transitions from marine to freshwater environments(Judd 2012; Hasan et al. 2017; Bonga 1973). Osmoregulation is fundamental during the early stages of freshwater adaptation(McCairns & Bernatchez 2009; Kusakabe et al. 2019; Divino et al. 2016). Adaptation to freshwater environments by marine populations is also characterized by marked behavioral changes mediated by structures in the brain(Park & Bell 2010; Bensky & Bell 2022). When threespine stickleback from a freshwater environment was exposed to predation risk, selection favored aggressive individuals(Bell & Sih 2007). There is also heritable differences in schooling behavior between marine and freshwater stickleback(Wark et al. 2011; Greenwood et al. 2013)

In this study we examined differentially expressed genes in the gill and brain of specimens collected from one marine and two freshwater populations of threespine stickleback fish in natural settings. One of the freshwater populations has only recently diverged from a marine progenitor, while the other is putatively a long-established freshwater population. We found differentially expressed genes between freshwater and marine populations in both brain and gills that may underlie certain molecular processes pertinent to freshwater adaptation, such as calcium regulation in the gill and fatty acid metabolism in the brain. Our results indicate that single nucleotide polymorphisms showing signatures of selection and rapid freshwater adaptation were enriched in differentially expressed genes. The SNPs that were enriched in differentially expressed genes mostly had low to moderate predicted effects on the resulting protein function or structure, perhaps indicating multiple variants acting in concert, but we also identified a high impact SNP that was associated with the expression of the *col8a1b* gene in a dosage dependent manner.

## Results

### Population sampling and demography

We collected brain and gill tissues from threespine stickleback individuals from three distinct Alaskan populations in the Anchorage/Matanuska-Susitna Valley region: Rabbit Slough, Cheney Lake, and Cornelius Lake (Methods, Fig. 1A). Cheney is approximately 60km south of Rabbit Slough, while Cornelius is 13km north of Rabbit Slough. We sampled five individuals from each population and generated TagSeq gene expression datasets from gill and brain tissues (total n=30, Methods). Rabbit Slough is an anadromous population that has been used to found many new freshwater populations including Cheney Lake(Bell et al. 2016; Aguirre et al. 2022). In 2009, 3,000 anadromous individuals from Rabbit Slough were transplanted to Cheney Lake in order to re-establish the stickleback population following their extirpation, which resulted from an effort to eliminate invasive northern pike (*Esox lucius*) from the lake.. Previous phenotypic and genomic studies revealed rapid adaptation to this new freshwater environment(Bell et al. 2016; Roberts Kingman et al. 2021; Wund et al. 2016).

**Fig. 1:**
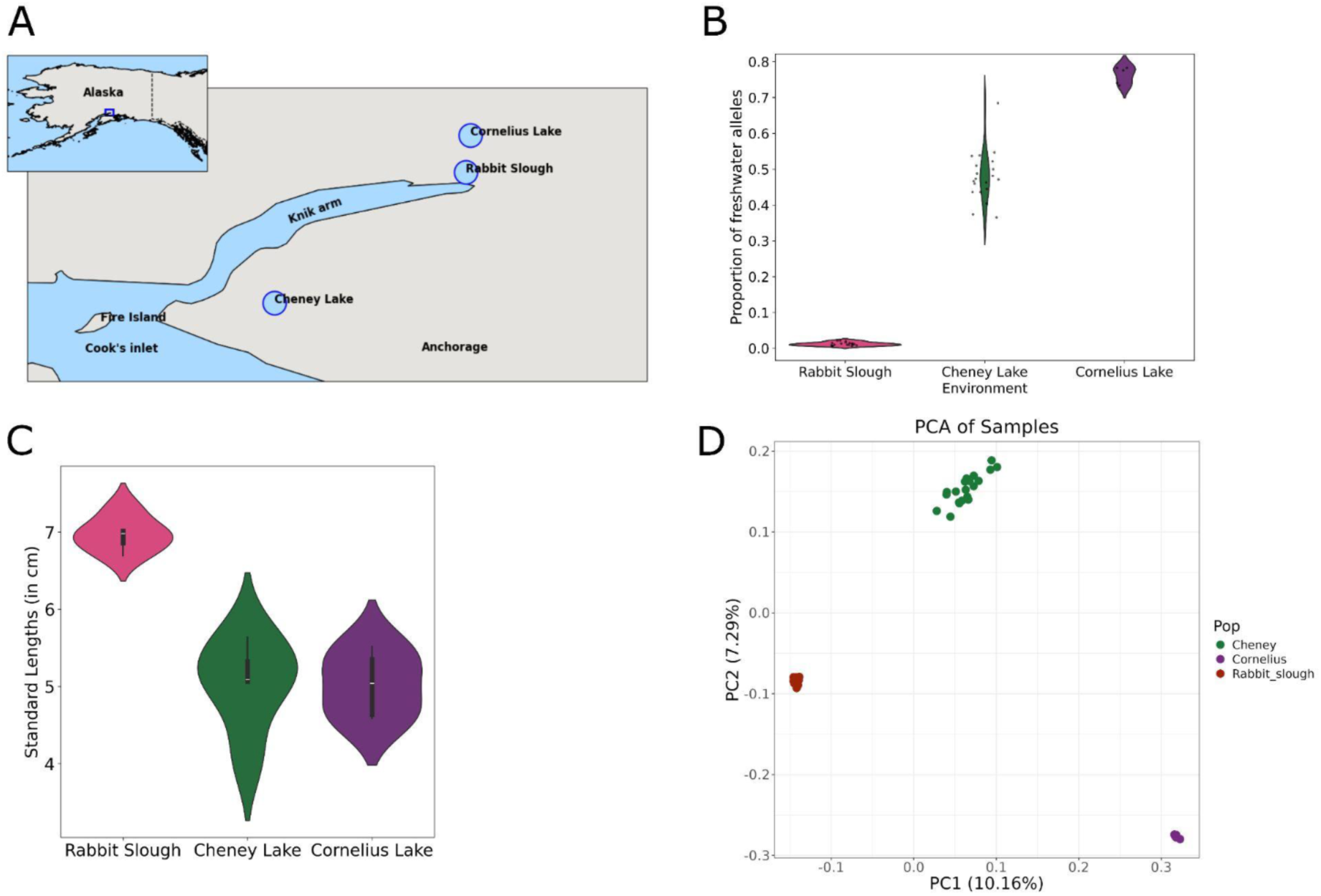
**A)** Location of lakes used in this study **B)** Proportion of freshwater adaptive alleles per individual for samples collected from the three environments **C)** Standard lengths of samples used for RNA sequencing **D)** Principal Components Analysis (PCA) using genome wide SNPs.

The threespine stickleback population found in Cornelius Lake has been putatively established for a much longer time than Cheney (assumed to be several thousand years); however, no previous study has determined the demographic history of this threespine stickleback population. Therefore, in order to explore its population genetic structure, we performed whole genome sequencing (WGS) for 5 Cornelius Lake specimens to approximately 6X coverage. We used an approximate genotype likelihood approach to call the diploid state of alleles at freshwater-adaptive loci previously identified by (Roberts Kingman et al. 2021) as homozygous oceanic, heterozygous, or homozygous freshwater. The genotypes at these loci were predominantly freshwater (Fig. S1). On average, each individual had ∼75% freshwater content, which reflects what is observed in other established freshwater populations in the Pacific Northwest (Fig. 2C, S2). In comparison, the average freshwater content in Rabbit Slough (based on n=20, 20x WGS published in (Roberts Kingman et al. 2021)) and Cheney Lake (based on n=20, 30x WGS published in (Kwakye et al. 2025)) was 1.2% and 48% respectively (Fig. 2C).

**Fig. 2:**
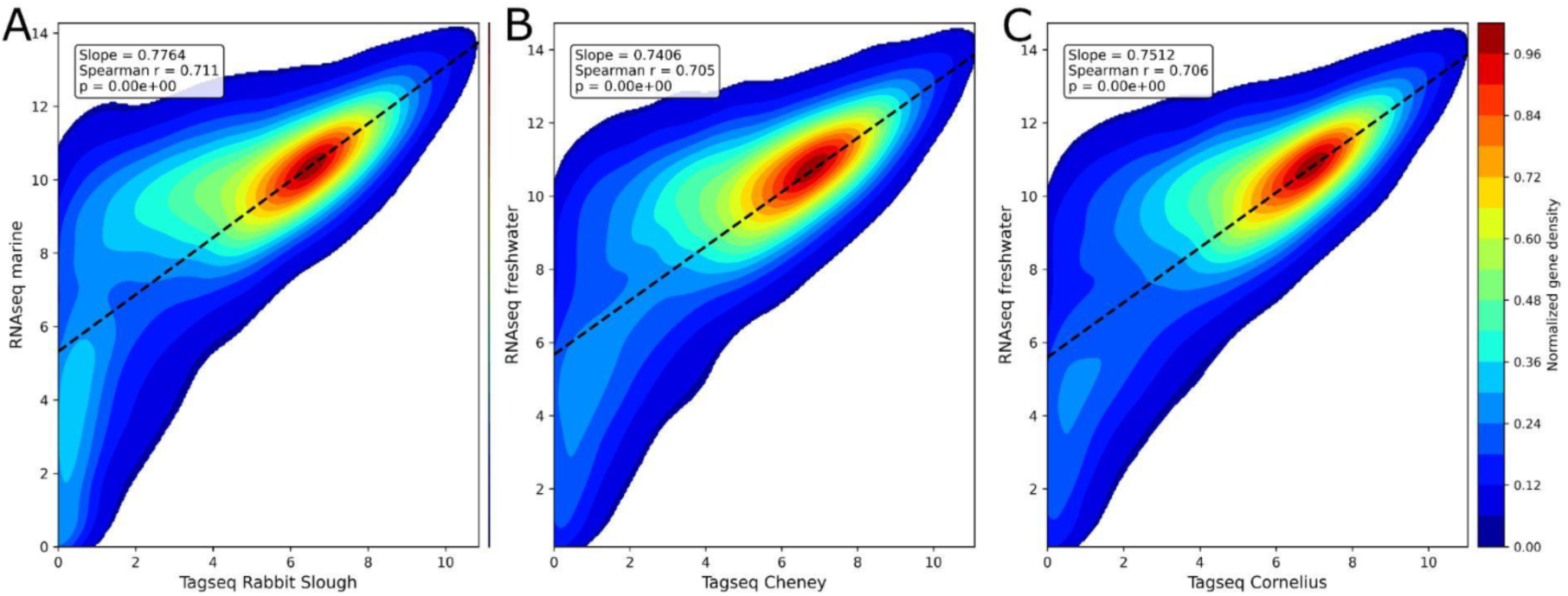
Comparison of bulk RNAseq dataset from Verta and Jones 2019 to TagSeq dataset in the present study. Correlation of gene counts estimated using TagSeq for **A)** Rabbit Slough individuals vs marine individuals collected from Little Campbell River **B)** Cheney individuals vs freshwater individuals collected from Little Campbell River. **C)** Cornelius individuals vs freshwater individuals collected from Little Campbell River.

The average standard length of Rabbit Slough specimens was 7cm, consistent with most marine threespine sticklebacks, whereas specimens from Cheney and Cornelius Lakes had standard lengths of 5.04cm and 5.02cm respectively, characteristic of mature freshwater sticklebacks(Baker 1994; DeFaveri & Merilä 2013) (Fig. 1B). We used Principal component analysis (PCA) and pairwise F_ST_ to determine the population genetic structure amongst the three populations used in this study. PC1 separated the samples based on population, with Cheney being intermediate between Rabbit Slough, from which it was recently derived, and Cornelius Lake, which is an established freshwater population. Pairwise *F_ST_* analyses showed similar patterns of population differentiation, with a mean genome-wide *F_ST_* of 0.033 between Rabbit Slough and Cheney Lake samples; 0.084 between Rabbit Slough and Cornelius Lake samples; and 0.07 between Cheney and Cornelius Lake samples.

When we considered only SNPs within regions of the genome that have alleles that consistently increase rapidly during freshwater adaptation (which we refer to as tempo SNPs in this study, see below), the *F_ST_* was 0.54 for Rabbit Slough versus Cheney Lake; 0.875 for Rabbit Slough vs Cornelius Lake; and 0.866 for Cheney versus Cornelius Lake. We also estimated the *F_ST_* after filtering out the tempo SNPs, leaving only putatively neutral sites. In this case the F_ST_ values were similar to those observed across the whole genome; the *F_ST_* between Rabbit Slough and Cheney was 0.032; Rabbit Slough vs Cornelius was 0.083; and Cheney vs Cornelius was 0.072. These results are consistent with the demographic and known or putative adaptive histories of these populations. Cheney Lake was recently founded using stickleback sampled from Rabbit Slough and thus these two populations are most similar genome-wide relative to Cornelius Lake, which likely diverged from anadromous ancestors much further in the past. However, these genome-wide differences driven by drift are accompanied with a substantial convergent freshwater adaptation that has occurred at both Cheney and Cornelius.

### Comparing gene expression from TagSeq and RNAseq

To determine whether our TagSeq approach for measuring gene expression could substantially bias downstream analyses, we compared the gene counts estimated from using TagSeq for gills to those estimated from bulk RNAseq for the same tissue from Verta and Jones 2019(Verta & Jones 2019). We processed the raw reads from the bulk RNAseq using a similar pipeline as Verta and Jones 2019(Verta & Jones 2019) (Methods).

We observed a very high correlation between the gene counts for the marine individuals from Rabbit Slough estimated using the TagSeq pipeline and the marine individuals from the bulk RNAseq (Fig. 2A; *r*=0.71, *p-value*<<0.001). Similarly, there was a high correlation between the freshwater samples (Cheney TagSeq vs freshwater RNAseq;*p*=0.00, *r*=0.705; Cornelius TagSeq vs freshwater RNAseq; *p*=0.00, *r*=0.706, Fig. 2B and 2C). We observed a weaker correlation when we compared expressed genes from the bulk RNAseq (from the gill tissue) to the expressed genes in the brain from Tagseq (Fig. S3). Notably, in all the comparisons, certain genes are expressed in the RNAseq dataset but are absent in the TagSeq, which is likely as a result of the lower coverage produced by TagSeq. Overall, the strong correlation observed across the comparisons demonstrates that expression levels measured by TagSeq are generally consistent with those measured by standard RNA-seq.

### Differentially expressed genes (DEGs) in the gill and brain

PCA of the normalized gene expression counts showed that the tissue of origin explained the majority of the variation (PC1=64.22%) among all the samples (Fig. 3A). After clustering by tissue, samples clustered on PC2 based primarily on population of origin (Rabbit Slough, Cheney or Cornelius).

**Fig. 3:**
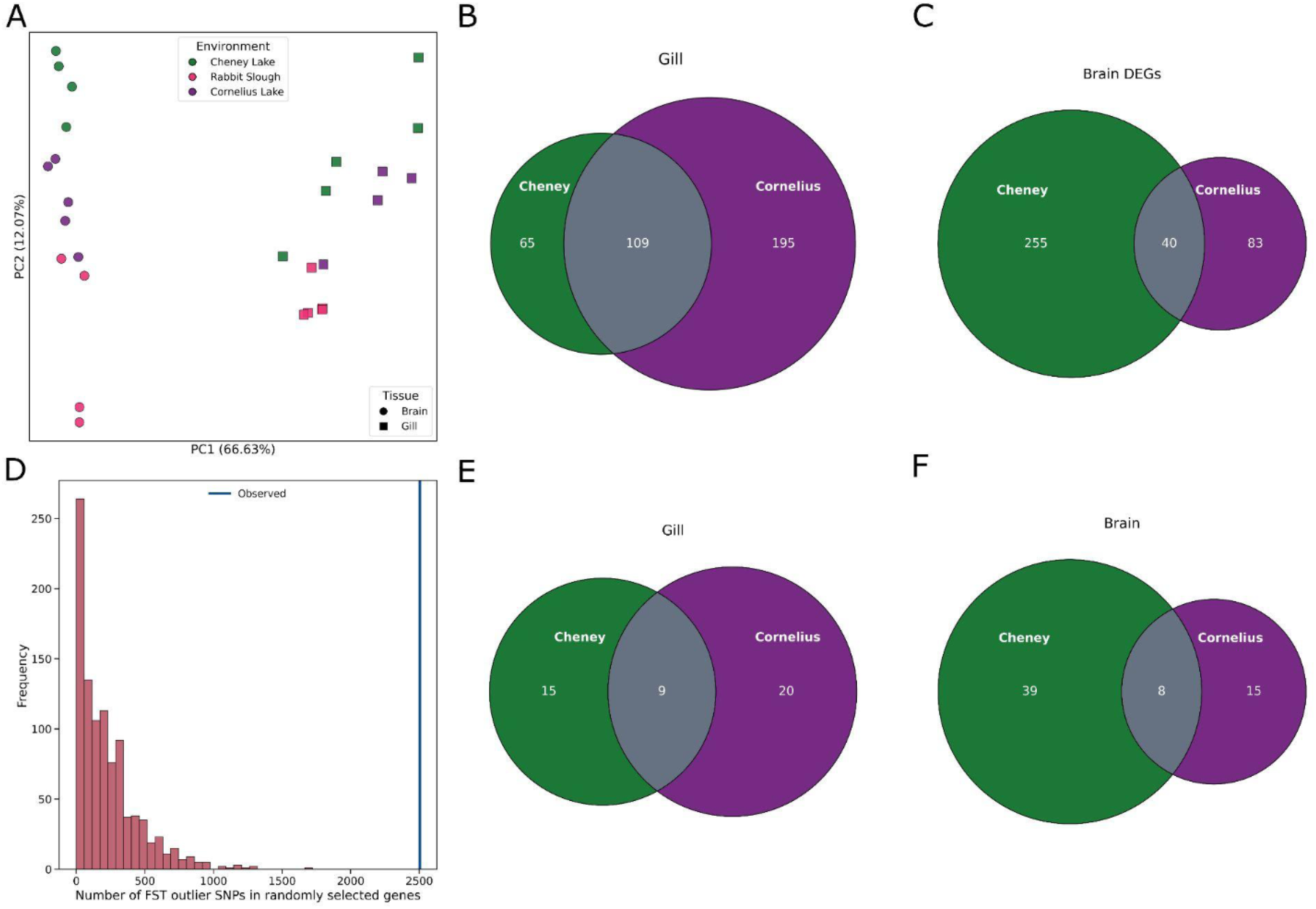
**A)** Principal Components Analyses (PCA) using gene expression profiles of all genes across all populations. Venn diagrams of differentially expressed genes (DEGs) when comparing the expression patterns in Cheney and Cornelius samples to Rabbit Slough for **B)** Gill and **C)** Brain. **D)** An example of our empirical approach for testing for enrichment of F_ST_ outliers and tempo SNPs in DEGs. Venn diagrams of the number of DEGs with F_ST_ outlier SNPs and tempo SNPs for **E)** Gill and **F)** Brain.

We detected expression of 14,187 genes in the gill in all samples across all three populations. Following normalization and correction for biological and technical variability using DESeq2(Love et al. 2014), we identified differentially expressed genes (DEGs) between environments i.e., Rabbit Slough (marine environment) vs Cheney and Cornelius Lakes (freshwater environments). The comparison between Rabbit Slough and Cheney Lake revealed 174 DEGs (log2FC > 2, adjusted *p-value* < 0.05) with the majority (n=159, 91%) being upregulated in Cheney Lake. There were 304 DEGs between Rabbit Slough and Cornelius Lake, of which 265 (87%) were upregulated in Cornelius Lake. There were 109 DEGs that were observed in both freshwater environments (Expected overlap: 3.65, Hypergeometric *p-value*: 8.557e-145, Fig. 3B), 105 (96%) of which were upregulated.

The brain had a lower number of total genes (12,290) expressed across the three populations. We found 295 DEGs between Rabbit Slough and Cheney Lake, with 266 (90%) being upregulated in Cheney Lake (log2FC > 2, adjusted *p-value* < 0.05). There were 123 DEGs found between Rabbit Slough and Cornelius Lake, with 88 (72%) being upregulated in Cornelius Lake (Fig. 3C). Thus, compared with Rabbit Slough samples, Cornelius Lake samples had more DEGs in the gill, while Cheney Lake samples had more DEGs in the brain. There were 40 DEGs that were observed in both freshwater environments (Expected overlap:2.93, Hypergeometric *p-value*: 5.270e-35, Fig. 3C). Among the 40 DEGs found in both freshwater environments, 34 were upregulated in the Cheney samples, while 32 were upregulated in the Cornelius samples. All the six DEGs that were downregulated in the Cheney samples were also downregulated in the Cornelius samples.

We aimed to determine whether the observed systematic upregulation of genes was a biological result of the pre–RNA extraction treatment or a technical artifact introduced by low-coverage RNA sequencing and bioinformatic pipelines. Many previous studies have measured gene expression using fish raised in common garden environments, while we measured gene expression in tissues extracted from wild-caught fish. Since the expression of housekeeping genes would remain unchanged across different environments, we can use their expression patterns to disentangle the possible cause of the observed upregulation of genes. We determined the expression of housekeeping genes from (Hibbeler et al. 2008), including: *actb1, gapdh, rpl13a, hprt1, tfb2m, taf2,* and compared them to our own results. In our study, the expression of all housekeeping genes remained unchanged in both marine and freshwater environments genes except *gapdh,* which was found to be differentially expressed in both Rabbit Slough vs Cheney Lake and Rabbit Slough vs Cornelius Lake comparisons (Fig. S4, S5). We also observed that the expression of these genes remained stable in the marine and freshwater samples reported in Verta and Jones 2019(Verta & Jones 2019). Though *gapdh* was not differentially expressed in their dataset there was an increase in its expression in freshwater individuals (Fig. S6). The observed differential expression of *gapdh* in our dataset likely reflects environmental stress encountered by wild-caught fish, consistent with previous studies that highlight the gene’s high expression variance in response to oxidative stressors(Camacho-Jiménez et al. 2023; Barber et al. 2005; Babington et al. 2025).

### Signatures of positive selection associated with DEGs

To determine if DEGs were associated with known freshwater adaptive loci, we examined to what extent the genes overlapped with approximately 33,000 SNPs associated with freshwater adaptation. These SNPs are found in 341 loci previously determined to have alleles that increase rapidly during freshwater adaptation in three recently established freshwater populations derived from an anadromous ancestor(Roberts Kingman et al. 2021). We refer to these SNPs as ‘tempo SNPs’. We also analyzed DEGs in relation to loci showing signatures of population differentiation by estimating the per-SNP *F_ST_* between each of the two freshwater populations and Rabbit Slough. For each DEG, we extracted SNPs found up to 10 Kbp upstream of its start site and 10 Kbp downstream of the end of the gene in order to capture putative *cis*-regulatory regions, including promoters and enhancers. We employed *F_ST_* outlier analyses to select SNPs underlying population differentiation. We set the *F_ST_* outlier threshold to 99th percentile to include only SNPs showing the strongest signals of genetic differentiation as such SNPs are more likely to be affected by divergent selection rather than demographic processes.

There were 63,651 *F_ST_* outlier SNPs found between Rabbit Slough and Cheney Lake samples, of which 546 SNPs were found in 24 gill DEGs between the same populations (Table S1). Out of these 546 SNPs, 10 of them were tempo SNPs, all of which were associated with the gene *pvalb4* (Table 1). For Rabbit Slough and Cornelius Lake samples, there were 100,077 *F_ST_* outlier SNPs, 2,503 of which were associated with 29 gill DEGs (Table S2). Out of these 2,503 F_ST_ outlier SNPs, 270 were tempo SNPs and were associated with 11 gill DEGs, including *pvalb4* (Table 1). Out of the 24 gill DEGs associated with *F_ST_* outlier SNPs between Rabbit Slough and Cheney Lake samples, nine of them overlapped the 29 gill DEGs associated with *F_ST_* outlier SNPs between Rabbit Slough and Cornelius Lake samples (Fig. 3E). This overlap was statistically significant (Hypergeometric p–value=2.0e-19).

**Table 1:**
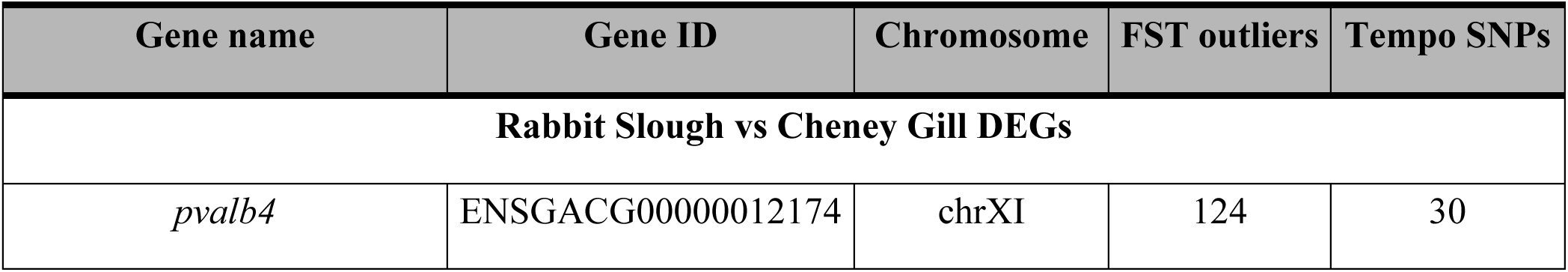
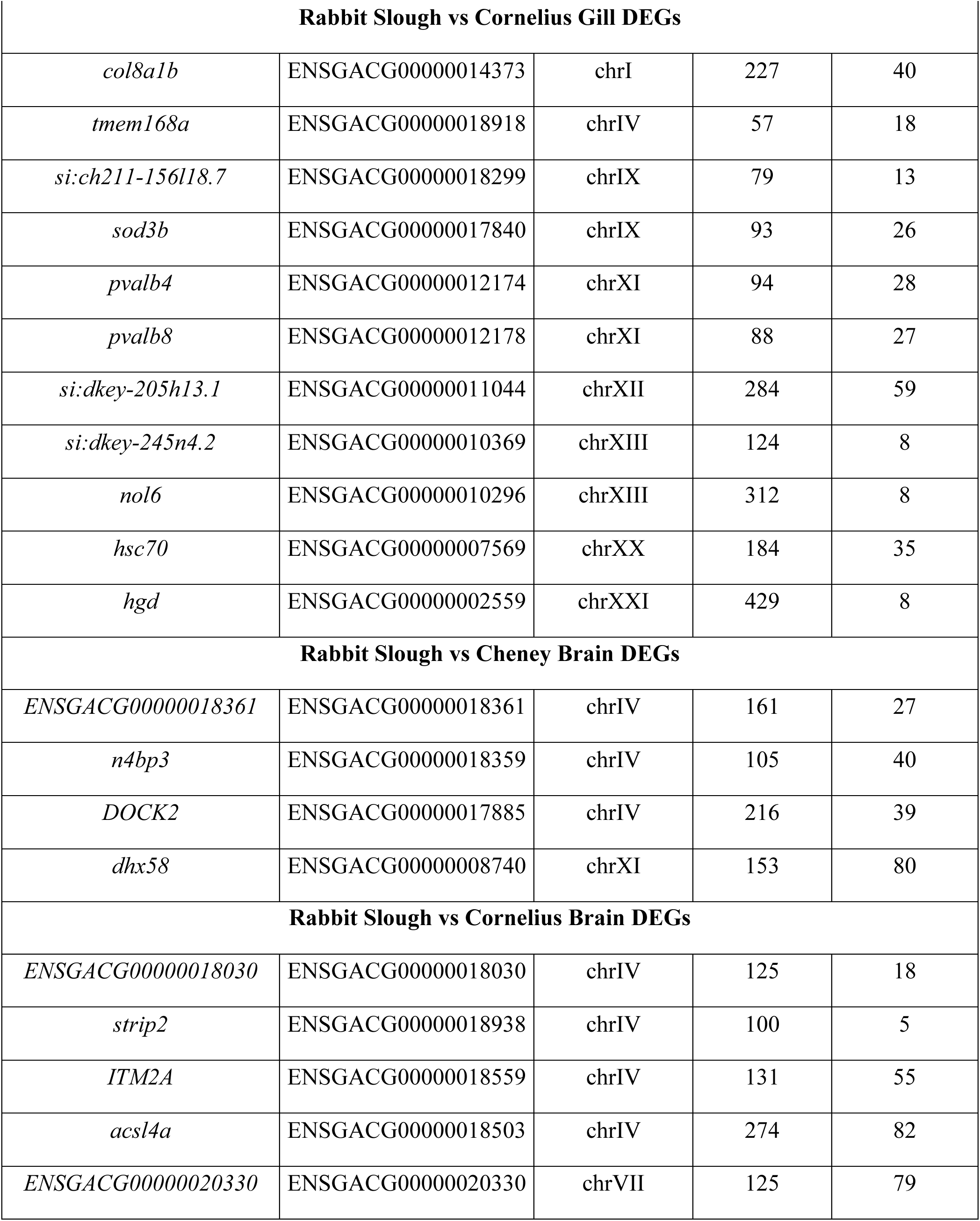
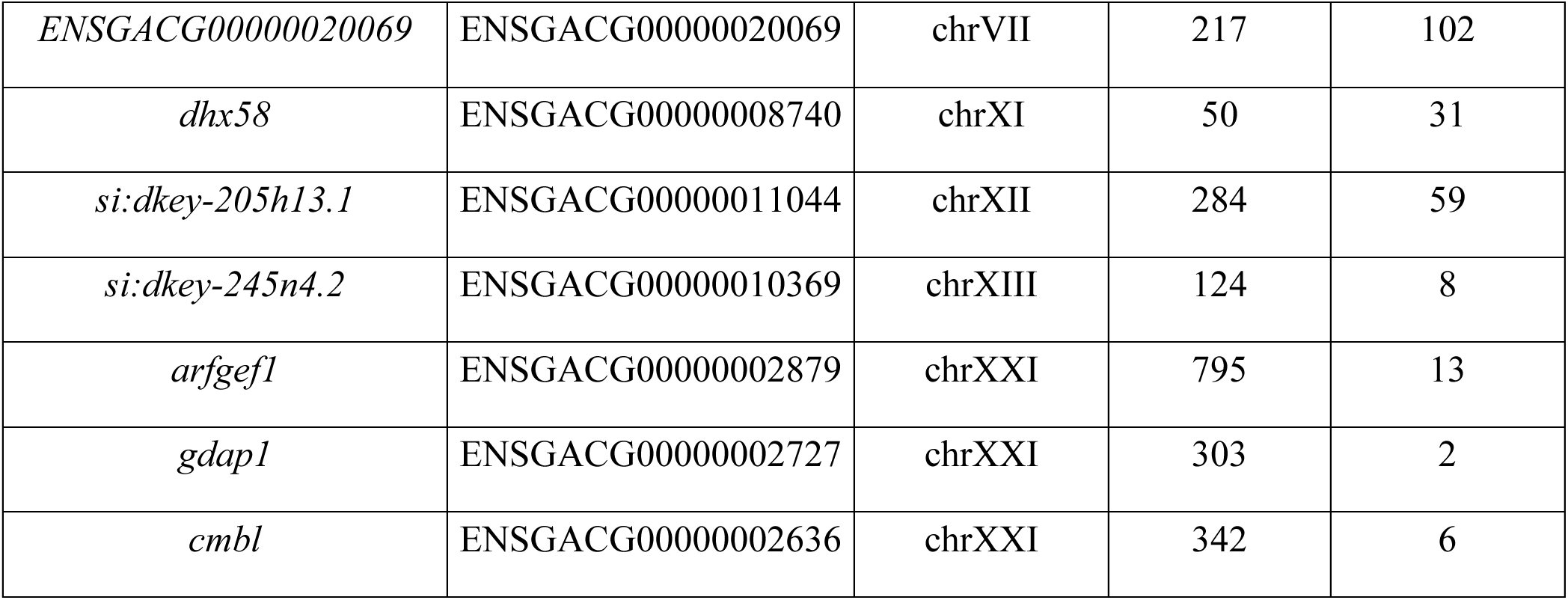
DEGs with both F_ST_ outliers and tempo SNPs.

There were 1144 *F_ST_* outlier SNPs found between Rabbit Slough and Cheney Lake samples that were associated with 47 of the brain DEGs (Table S3). Of these, 186 were tempo SNPs and were associated with four DEGs (Table 1). There were 3,381 *F_ST_* outlier SNPs found between Rabbit Slough and Cornelius samples that were associated with 23 brain DEGs (Table S4). Of these, 460 were tempo SNPs and were associated with 12 DEGs, including the gene, *dhx58*, which was among the four genes also found in the Rabbit Slough and Cheney comparison (Table 1). Out of the 47 brain DEGs harboring *F_ST_* outlier SNPs between Rabbit Slough and Cheney Lake samples, eight of them were among the 23 brain DEGs with *F_ST_* outlier SNPs between Rabbit Slough and Cornelius samples (Fig. 3F, Hypergeometric test p–value=1.1e-14).

### Enrichment of signatures of selection in DEGs

We then tested whether the *F_ST_* outlier SNPs and tempo SNPs associated with DEGs were significantly enriched compared to the rest of the genome using the following approach. For each set of DEGs, we chose an equal number of genes from the rest of the genome that match the total length of the DEG gene set, with a tolerance of ± 1Kb. We generated 1000 random gene sets matching this length criterion, and estimated the number of *F_ST_* outlier SNPs and tempo SNPs within each of them, creating an empirical null distribution for comparison to our observed number of F_ST_ outlier SNPs and tempo SNPs in each DEG set (Fig. 3D).

We observed that *F_ST_* outlier SNPs were significantly enriched in the DEGs from the comparison between Rabbit Slough and Cornelius Lake samples for both gill and brain tissues (p–value=0.0 in both cases). However, we did not find statistically significant enrichment for *F_ST_* outlier SNPs in DEGs from comparison between Rabbit Slough and Cheney Lake samples in these same tissues (gill *p–value*=0.056, brain *p–value*=0.106). Similar to the *F_ST_* outliers, the tempo SNPs were also significantly enriched in the gill DEGs (*p–value*=0.016) and brain DEGs (*p–value*=0.0) for the comparison between the Rabbit Slough and Cornelius samples, while for the Rabbit Slough versus Cheney comparison, the tempo SNPs were only significantly enriched in the brain (*p–value*=0.012) but not the gill (*p–value*=0.054). Overall, these results suggest that signatures of population differentiation and signatures of rapid freshwater adaptation were enriched in DEGs compared to the rest of the genome, but particularly in the more longstanding Cornelius population.

### Parvalbumin expression in the gill

Considering that the *pvalb4* gene was differentially upregulated in the gill in both freshwater environments (Fig. 4A) as well as harboring substantial *F_ST_* outlier SNPs and tempo SNPs, we explored its expression in the gill RNAseq dataset from Verta and Jones 2019(Verta & Jones 2019). Based on the criteria we used in identifying differentially expressed genes in the TagSeq dataset, *pvalb4* was not significantly differentially expressed between the marine and freshwater samples from Verta and Jones 2019 (log2FoldChange= –0.107, adjusted *p-value*=0.980). However, when we considered the expression of this gene compared to the rest of the genes in the genome, *pvalb4* was among the most highly expressed genes in the genome (Fig. 4B), suggesting that it may be involved in one or more key functions in threespine stickleback. The gene also showed elevated expression in females compared to males in this dataset (Fig. 4C), although this trend was not replicated in the TagSeq dataset, where males instead showed elevated expression.

**Fig. 4:**
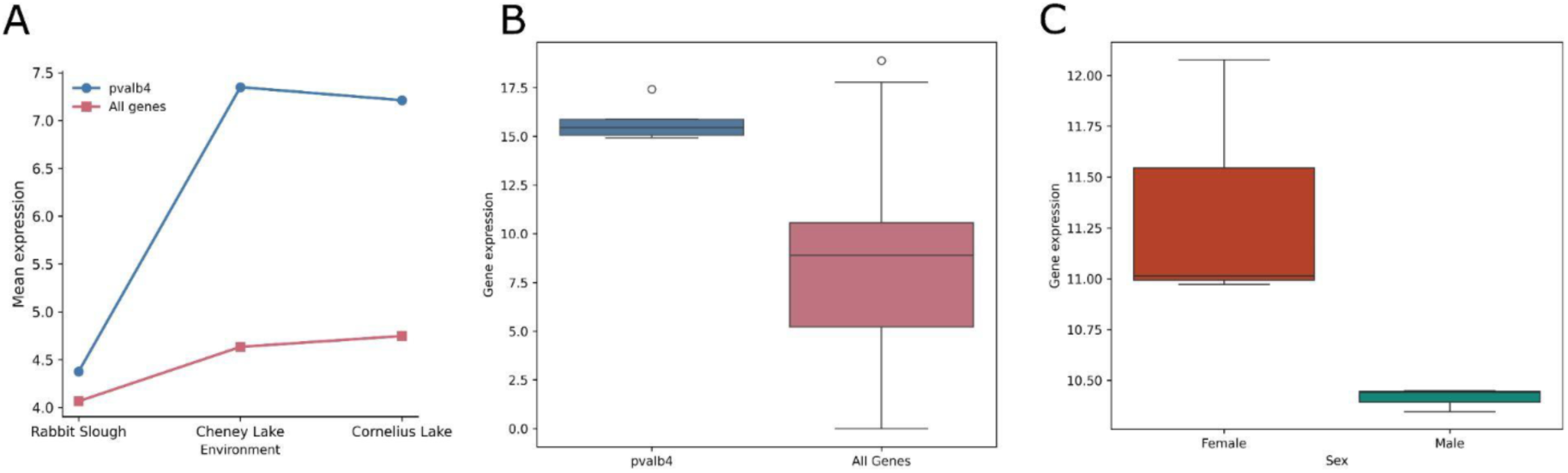
A) Expression of parvalbumin(*pvalb4)* compared to all other genes in the gill of Rabbit Slough, Cheney and Cornelius Lakes. B) Expression of *pvalb4* gene in the gill in samples from Verta and Jones 2019 in comparison to other expression levels of other genes in the genome. C) Expression of *pvalb4* gene in the gill samples from Verta and Jones 2019 between females (n=3) and males (n=3).

### Genetic load in DEGs

We next aimed to determine the predicted effects of freshwater/marine-divergent SNPs associated with DEGs on protein function and structure. In this analysis, we focused on tempo SNPs that were also *F_ST_* outliers for all sets of DEGs, and referred to these SNPs as ‘candidate SNPs’. In total, there were 946 candidate SNPs across all four DEG sets. We annotated candidate SNPs based on their predicted effects on protein function using snpEff(Janssen et al. 2023; Cingolani et al. 2012), which assigns SNPs to impact classes: low, moderate, high and modifier. We also determined the evolutionary constraint on candidate SNPs using their site-specific GERP scores(Davydov et al. 2010; Cooper et al. 2010)(Methods). SnpEff can assign multiple effects to the same variant. For example, if a gene has multiple transcripts, the program assigns effects to all transcripts; or if a variant is located upstream of one gene and downstream of another, it reports effect of the variant on both genes. We therefore report all effects assigned to each SNP.

We observed that most candidate SNPs had either low, moderate or modifier effects, with the majority being modifiers (Fig. 5A). The modifier subclass was predominantly used to annotate upstream or downstream variants, suggesting that freshwater adaptation is mostly mediated by *cis*-regulatory changes (Fig. 5B). The moderate effect variants were predominantly missense variants; 8 missense variants in Rabbit Slough vs Cheney Lake brain DEGs, 22 in Rabbit Slough vs Cornelius Lake brain DEGs and 31 in Rabbit Slough vs Cornelius Lake gill DEGs (Fig. 5C). The eight missense variants in the Rabbit Slough vs Cheney brain DEGs were distributed across three DEGs, with *dhx58* harboring three of them. The 22 missense variants in the Rabbit Slough vs Cornelius Lake brain DEGs were found across 8 DEGs, including five in the gene *acsl4a*, five in the gene *gpr174,* and one in *dhx58* that was also identified in the Rabbit Slough vs Cheney brain DEGs. There were 13 of the missense variants from the Rabbit Slough vs Cornelius Lake gill DEGs that were found in the gene *col8a1b,* while three were found in the gene *sod3b* and two in the gene *DHX15* (Table S5).

**Fig. 5:**
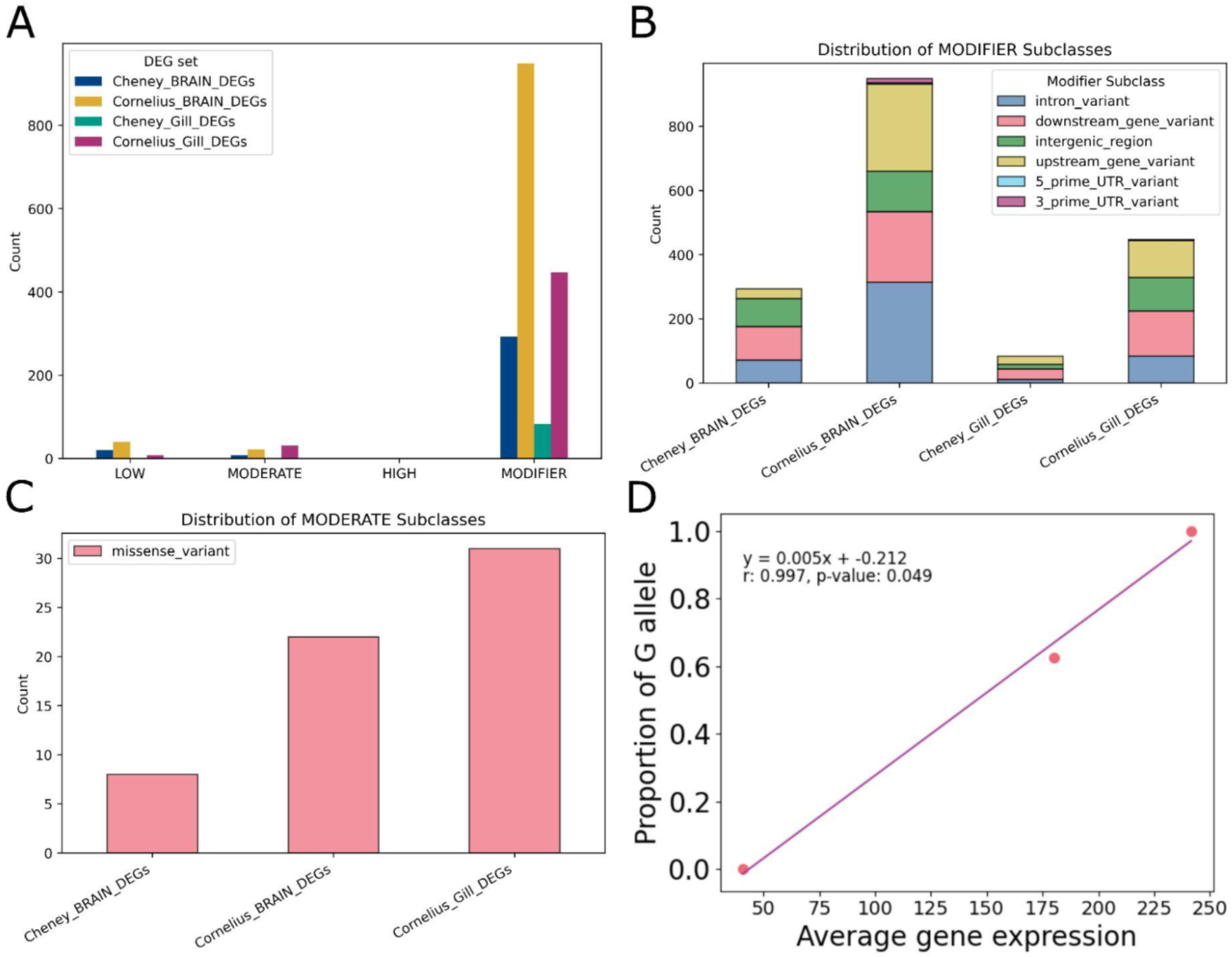
Predicted molecular effects of candidate SNPs. **A)** Distribution of molecular effects of 946 candidate SNPs as predicted by snpEff. Breakdown of the various components that make up the **B)** modifier impact class **C)** moderate impact class (all of which were missense variants). **D)** Correlation between the average gene expression in the gill tissue and the proportion of G alleles at a SNP((chrI:26106268 G > T) predicted to have a high impact on the gene *col8a1b*.

We identified a candidate SNP in the Rabbit Slough vs Cornelius Lake gill DEGs on chromosome I (chrI:26106268 G > T) with a putatively high-impact on the molecular function and/or structure of the *col8a1b* gene product. This SNP potentially leads to a premature stop codon c.1838C>A (p.Ser613*). The expression of the gene decreased in the direction of the T allele, which putatively leads to a stop mutation in place of a codon for serine. The expression of the *col8a1b* in the gill was substantially correlated with the allele counts in the three populations(Fig. 5D, *r*=0.98, *p*=0.122). All five individuals in the Cornelius sample were homozygous for the G allele, while the 20 Rabbit Slough individuals were homozygous for the T allele, with the observed heterozygosity in the Cheney population being 0.65(expected heterozygosity=0.30). The expression of the gene *col8a1b* in the brain was reduced compared to its expression in the gill(Fig. 5D, S10). The SNP is also found upstream of the gene *jagn1b* and is predicted to be in the modifier class for this gene, however, this gene was not expressed in either tissue.

Out of the 946 candidate SNPs, we retrieved the GERP scores for 139 of them. Out of these, 115 (83%) had GERP scores above 0. However, the distribution of GERP scores in sticklebacks is right-skewed, with a median (50th percentile) value of 7.55; 90th, 95th and 99th percentiles of 16, 17.20 and 19.90 respectively (Fig. S7). Out of the 139 candidate SNPs with GERP scores, 68 were found in the 50th percentile, 18 in the 90th percentile, 5 in the 95th percentile and 0 in the 99th percentile. Out of the five in the 95th percentile, four were found in Rabbit Slough and Cornelius brain DEGs (all of which were associated with the gene ENSGACG00000020069) and one in a Rabbit Slough and Cornelius gill DEG, which was associated with the *hgd* gene. All the SNPs had one allele fixed in the Rabbit Slough sample, while the alternate allele was fixed in the Cornelius sample.

We also determined the evolutionary constraints on specific DEGs harboring substantial *F_ST_* outlier SNPs and tempo SNPs such as *pvalb4* and missense variants such as *dhx58* and *col8a1b.* For sites where we retrieved the GERP scores in *pvalb4* and *dhx58*, we observed that all of the GERP scores were below the genome-wide median (Fig. S8 and S9). There were no sites with GERP scores in *col8a1b*.

### Gene ontology

To identify molecular processes that are enriched in each set of DEGs, we performed gene ontology (GO) analyses using Gprofiler(Kolberg et al. 2023). The GO term voltage-gated calcium channel complex was significantly enriched in the DEGs found in the Rabbit Slough vs Cheney Lake gill comparison, while distal renal tubular acidosis was significantly enriched in the DEGs in the Rabbit Slough vs Cheney Lake brain comparison. There were no enriched GOs in the DEGs for both tissues in the Rabbit Slough and Cornelius Lake comparisons.

## Discussion

### Differentially expressed genes in brain and gill

In this study we leveraged the repeated, rapid adaptation observed in threespine stickleback to explore which genomic signatures of selection likely influence gene expression changes, and to examine the mechanisms underlying how these signatures may shape expression patterns. We observed that signatures of selection were enriched in differentially expressed genes in the brain and gill tissues of samples collected from stickleback populations inhabiting different freshwater environments compared to the rest of the genome. The enriched signatures of selection were mostly located in regulatory regions of genes, consistent with previous studies(Jones et al. 2012; Verta & Jones 2019). However, we did find strong evidence for coding change at a gene leading to a premature stop codon that likely influences protein function in a dosage-dependent manner in the gill.

The number of expressed genes we identified in the gill is slightly less than Rodríguez-Ramírez and Peichel 2025(Rodríguez-Ramírez & Peichel 2025) (14,500 in the present study vs 16,688 genes from Rodríguez-Ramírez and Peichel 2025) who re-analyzed two previously published gene expression datasets from (Gibbons et al. 2017) and (Verta & Jones 2019). The differentially expressed genes we identified in the gill were about a third of the number identified by Rodríguez-Ramírez and Peichel 2025. The difference in expressed genes between the two studies likely result from the different sequencing approaches used by different studies. We utilized TagSeq, which involves sequencing the 3’ end of mRNAs, while the other studies re-analyzed by Rodríguez-Ramírez and Peichel 2025(Rodríguez-Ramírez & Peichel 2025) utilized bulk RNAseq with higher coverage.

Although Robertson et al. 2016(Robertson et al. 2016) did not compare marine and freshwater stickleback, the authors monitored the expression of immune specific genes including genes characteristic of innate immunity, immunosuppression, and helper T lymphocytes in stickleback from wild populations and lab raised fish. They observed that wild fish differed in the expression of immune specific genes characteristic of innate immunity, immunosuppression, and helper T lymphocytes (except the Th1-subtype of helper T cells) compared to lab raised fish. Importantly, wild fish had increased expression of all the immune specific genes. The differences in the patterns of gene expression in immune specific genes between wild caught fish and lab raised fish indicate that possibly the difference in the number of differentially expressed genes between this study and the previous studies was due to treatment of the samples prior to RNA extraction. In the studies that Rodríguez-Ramírez and Peichel 2025 re-analyzed, the fish were captured from the wild and reared in the lab for a period before extracting RNA, which, by design, depletes some variation. In the current study, we used RNA from tissues collected directly from wild-caught fish, which may be more variable, reducing power to detect genes with only subtle expression changes.

### Genes involved in molecular processes in gill and brain

Our focus on the brain and gill was based on their roles in osmoregulation and behavior during transitions from marine to freshwater environments(Judd 2012; Hasan et al. 2017; Bonga 1973; Bell & Sih 2007). In the gill, we found genes such as *tnnt1, pvalb4* and *atp6v1e1a* to be differentially expressed in both freshwater environments, relative to their anadromous ancestor, and also enriched for signatures of selection. *Tnnt1* encodes a subunit of troponin that binds to tropomyosin(Pato et al. 1981; Wei & Jin 2016). Through its role in anchoring the troponin complex to the thin filament, *Tnnt1* contributes to calcium-dependent regulation of gill muscle contraction, which may facilitate sustained ventilation and gaseous exchange across the gill filaments. Although muscle contraction depends on intracellular calcium release rather than environmental calcium directly, the reduced availability of external calcium in freshwater may impose physiological constraints that select for altered regulation of calcium-sensitive contractile proteins, including components of the troponin complex, in gill-associated muscles. In addition to *tnnt1*, *pvalb4*, which encodes the parvalbumin protein that chelates calcium, thereby contributing to calcium regulation(Heizmann 1984), was also differentially expressed in the gill. The gene *atp6v1e1a* encodes for a subunit of an ATP-dependent intracellular protein pump, which potentially is involved in influx of calcium into the gill. Although *cacnb1* was found to be differentially expressed only in Cheney, it encodes for a subunit of voltage-gated calcium channel that was found to be differentially expressed during a salinity change(Taugbøl et al. 2022).

*Pvalb4* was differentially upregulated in both freshwater environments and was enriched with *F_ST_* outliers and tempo SNPs. Parvalbumin has been implicated in marine-freshwater divergence in at least one other study, which found it to be differentially downregulated in the gill of freshwater stickleback(Verta & Jones 2019). Our re-analysis of a subset of the dataset from Verta and Jones 2019(Verta & Jones 2019) also indicated that the gene is highly expressed in sticklebacks in both marine and freshwater environments; however, according to our differential expression criteria, it was not classified as differentially expressed. Verta & Jones 2019 subjected wild-caught to brackish water (3.5 parts per thousand salinity) in the lab, which may mask the freshwater-specific upregulation of the *pvalb4* gene we observed in our wild-caught specimens. Parvalbumin has been shown to be involved in fast muscle contraction(Celio & Heizmann 1982), which may explain why it is highly expressed in a fish species with fast swimming activities(Zhang et al. 2017). In individuals of the *Xenopus allofraseri* that exhibit high levels of locomotion, *pvalb4* was found to be differentially expressed compared to those with reduced mobility(Ducret et al. 2021). Parvalbumin has also been linked to sustained locomotion in fishes(Seebacher & Walter 2012). These findings suggest that *pvalb4* may play key roles during the adaptation of threespine stickleback to freshwater environments, possibly by allowing the fish to escape predators. However, we note that the effect of an increase in *pvalb4* expression in anti-predation may be well tested with gene expression experiments that include other tissues, particularly the muscle tissue.

In threespine stickleback, the comparatively lower calcium concentration in freshwater environments has usually been suggested as a selective force driving lateral plate polymorphisms between the contrasting environments(Giles 1983; Smith et al. 2014) (but see (MacColl & Aucott 2014; Reimchen 1983)). Our results suggest that the contrasting calcium concentrations between environments may be driving phenotypic changes in the gill, perhaps more so than in the lateral plates. Our Gene Ontology analyses of the differentially expressed genes in the gill also suggested that voltage-gated calcium channel complex was significantly enriched in the Rabbit Slough vs Cheney gill DEGs. The enrichment of both tempo SNPs and *F_ST_* outliers in differentially expressed genes such as *pvalb4* may indicate that calcium concentration has been a selective force driving freshwater adaptation for a long time.

In the brain, genes such as *strip2*, *dhx58 and acsl4a* were found to be enriched with signatures of selection. *Acsl4a* encodes for a ligase that converts long-chain fatty acids into fatty acyl-CoA esters. In freshwater environments, the diet of threespine stickleback is deficient in long-chain unsaturated fatty acids such as docosahexaenoic acid (DHA), which are abundant in marine environments. Freshwater populations therefore have to rely on cellular biosynthesis of these fatty acids for normal growth and development. *Acsl4a* was previously found to be involved in lipogenesis(Lopes-Marques et al. 2013) in cavefish, suggesting that it may play key roles in fatty acid metabolism in stickleback. In sticklebacks, Ishikawa et al. 2019(Ishikawa et al. 2019) showed that *fads2*, a gene involved in DHA biosynthesis, is crucial for the colonization of new freshwater environments. *Fads2* is the rate-limiting enzyme in the poly-unsaturated fatty acid biosynthesis pathway and *Acsl4a* acts downstream of this enzyme. It remains unclear how the poly-unsaturated fatty acid biosynthesis pathway is implicated in the brain and the specific selective pressure involved but it is possible that this pathway may be involved in neurogenesis, synaptic function or inflammation(Bazinet & Layé 2014).

Inflammation in the brain may explain the differential expression of *dhx58* in the brain. This gene encodes an ATP-dependent RNA helicase that regulates type I interferon and is involved in antiviral innate immunity. In threespine stickleback, it is not only parasites such as the cestode, *Schistocephalus solidus*, that is ubiquitous in the organism’s habitat, but it may also co-evolve with viruses such as *Picornaviridae*, a family of nonenveloped RNA viruses(Hahn & Dheilly 2019). The presence of viruses may explain why genes involved in antiviral immunoregulation such as *dhx58* are differentially expressed. In two Alaskan populations with varying degrees of infection, Wohlleben et al. 2024(Wohlleben et al. 2024) found *dhx58* expression to be associated with *S. solidus* infection, although type I interferon which is regulated by the RNA helicase *dhx58* is involved in antiviral immunity.

### Enrichment of SNPs in differentially expressed genes

Our results showed that SNPs showing signatures of population differentiation and rapid freshwater adaptation were mostly enriched in DEGs compared to the rest of the genome. In Cheney Lake individuals, *F_ST_* outliers were not enriched in both brain and gill DEGs likely because the population recently diverged from Rabbit Slough stickleback. In the more putatively established freshwater population in Cornelius Lake, we observed enrichment of *F_ST_* outliers in DEGs from both tissues. Tempo SNPs, which are sites with signatures of rapid freshwater adaptation, were enriched in both tissues in Cornelius Lake fish and in the brain DEGs of Cheney Lake fish.

We found a SNP with putatively high impact on protein structure and function of *col8a1b* gene which codes for the alpha-1 chain of the collagen VIII protein. A transcript of the gene *col8a1b* gene is found on the reverse strand of chromosome I and has three exons but the start codon is found on exon 2 which means that the first exon is primarily located in the 5’ UTR. The SNP is located in the third exon of *col8a1b* where it results in a stop codon at c.1838C>A, truncating the 746-amino-acid protein at position 613. The truncated protein may be dysfunctional because collagen VIII assembly requires the truncated carboxyl-terminus domain to initiate trimerization<u>(Boudko et al. 2009)</u>. The stop codon may be acting as a cis-eQTL, influencing *col8a1b* expression in a dosage-dependent manner; the T(A on the reverse strand) allele is associated with significantly reduced transcript levels, a pattern consistent with incomplete dominance. The SNP deviated substantially from Hardy-Weinberg equilibrium, which could possibly indicate substantial selection pressure. In zebrafish, this gene is involved in vertebral column segmentation during development(Gray et al. 2014).

Aside from this SNP, we observed that the majority of the signatures of selection were located in *cis-*regulatory regions, and included variants up- and down-stream of genes, as well as synonymous variants consistent with previous studies(Jones et al. 2012; Verta & Jones 2019). There were also missense variants in genes such as *dhx58, acsl4a, gpr174, col8a1b, sod3b and DHX15.* We have described the molecular functions of the majority of these genes above, except *gpr174, sod3b* and *DHX15. DHX15* is an RNA helicase that can function as a viral sensor in the innate immune system(Xing et al. 2021; Pattabhi et al. 2019) and may be required for antiviral immunoregulation during freshwater adaptation. *Sod3b* is involved in the regulation of oxidative stress during adaptation to extreme conditions in an Antarctic blackfin icefish(Kim et al. 2019).

Our analyses of the evolutionary constraints on single nucleotide polymorphisms found in differentially expressed genes showed that genes such as *hgd* and ENSGACG00000020069 harbor highly-conserved SNPs. In contrast, genes that harbor substantial numbers of SNPs showing signs of population differentiation and rapid freshwater adaptation such as *pvalb4* and those with missense variants such as *dhx58* were not highly conserved. We however note that a previous study found copy number variation near the transcription start site of ENSGACG00000020069 where the marine and freshwater copy numbers are different(Lowe et al. 2018). ENSGACG00000020069 produces a c-c motif chemokine protein, which is part of a family of signalling proteins involved in the chemotaxis of immune cells.

### Upregulation of genes during freshwater adaptation

Our analyses showed that the vast majority of the genes that were differentially expressed in both tissues were upregulated in the freshwater environments. Mommer and Bell 2014(Mommer & Bell 2014) measured the differential gene expression of embryos from mothers exposed to predation risk and compared them to embryos from control mothers (no exposure to predators). The authors found that 66% of the differentially expressed genes were upregulated in the embryos from the predation-risk mothers. This study measured differential gene expression using the whole embryo, therefore their results do not reflect tissue specific gene expression changes. Thus, the increased proportion of upregulated genes that we observed may be systemic, rather than tissue specific.

In Gibbons et al. 2017(Gibbons et al. 2017), the authors subjected marine and freshwater stickleback fish to varying salinities and measured the patterns of differential gene expression in the gill tissue. When they compared gene expression in the gill of fish reared in 0 ppt and 30 ppt, they observed almost equal proportions of upregulated genes to downregulated genes. They observed similar patterns when they compared marine and freshwater ecotypes. Verta and Jones 2019(Verta & Jones 2019) also treated marine and freshwater ecotypes to varying degrees of salinities and measured the differential expression of genes in the gill. Although the authors did not provide exact counts of up- and downregulated genes, our reanalysis indicated that the numbers were roughly equal. It may be that studies that measured gene expression using threespine stickleback raised in common garden environments eliminate certain triggers–such as pathogens, or predators in the natural setting that cause certain genes to turn on. On the other hand, quantifying gene expression from wild-caught fish might capture expression patterns induced by environmental variables that turn on expression of genes. For instance, *gapdh* may be upregulated due to an increased demand for ATP. In addition, Verta and Jones 2019(Verta & Jones 2019) suggested that differentially expressed genes found in regions of marine–freshwater divergence tend to be predominantly upregulated, which is in line with our finding differentially expressed genes, most of which were upregulated, were also enriched with signatures of population differentiation and rapid freshwater adaptation.

Our findings suggest that threespine stickleback in freshwater environments may mount a systematic upregulation of genes associated with critical physiological processes, such as calcium regulation and fatty acid metabolism. Our analysis revealed that *gapdh,* a metabolic gene that is frequently used as a housekeeping gene, was differentially expressed in the freshwater populations. This shift in expression may reflect the elevated metabolic costs of active ion-regulation required to maintain homeostasis in ion-poor environments. Thus, increased metabolic and physiological demands appear to drive a broad transcriptomic shift toward increased expression of essential survival genes in freshwater habitats. Possibly, DNA hypermethylation silences the expression of these genes in the marine environment and the change in environment to freshwater induces genome-wide hypomethylation(Hu & Barrett 2023), which then increases the expression of genes by making them more accessible for transcription. Genome-wide hypomethylation is however inconsistent with studies that identified hypermethylation in both marine and freshwater samples(Smith et al. 2015). It is also possible for chromatin remodelling and accessibility to mediate such widespread upregulation of genes when marine stickleback encounter a new freshwater environment. Further studies are needed to better understand the role of gene upregulation in freshwater adaptation.

## Conclusion

Recent advancements in genomic scans have revealed many signatures of selection. Yet it remains unclear how these signatures of selection mediate phenotypic changes. In this study, We showed that single nucleotide polymorphisms that underlie freshwater adaptation are enriched in differentially expressed genes. These SNPs were predominantly found in regulatory regions and likely have small effects on protein function and structure. The SNPs with predicted high-impact on protein function and structure predominantly showed dosage-dependent effect on the gene expression. Together, the results add to growing efforts to connect alleles identified by more than two decades of genome scans to the phenotypes they might influence.

## Materials and Methods

### Sampling of threespine stickleback, RNA preservation and extraction

Threespine stickleback were collected between June 2 and June 10 2023 from Cheney Lake (61.204, –149.76), Cornelius Lake (61.628, -149.255) and Rabbit Slough (61.532, –149.266) in Alaska, USA. Minnow traps were deployed to collect stickleback samples from these three locations. Five traps were set at Cheney Lake on the west arm of the Northern peninsula of the lake at 4:45pm on June 2, 2023 (air temperature 10.5℃, water temperature 12℃), yielding 30 sticklebacks upon recovery on June 3 at 2:32 PM. On June 3, ten traps were deployed at Rabbit Slough at the stream flow near the Glenn Highway at 12:00 PM (air temperature 18℃, water temperature 13℃). We recovered the traps on June 4 at 12:20 PM, yielding 431 fish. Ten traps were also set at Cornelius Lake, near North Engstrom Road at 1:30 PM on June 3 (air temperature: 16.5℃, water temperature 12.5℃) and recovered on June 4 at 11:00 AM, yielding 375 fish. For all the three locations, a subset of captured individuals (total 13 for Cheney Lake, 41 for Rabbit Slough, and 47 for Cornelius Lake) were kept. We sacrificed five to 10 individuals shortly after collection using MS-222, and then harvested the brain and gill tissues. The tissues were immediately preserved in RNAlater and ultimately stored at -80℃ for subsequent molecular analysis. All procedures were approved by The College of New Jersey Institutional Care and Use of Animals Committee (protocols 1908-001MW1A1 and 2002-001MW1A3).

### Measurement of standard lengths

For each fish from which we harvested brain and gill tissues, we took a photo and estimated the standard length (SL) using imageJ software from the photos. The SL is measured as the distance from the anterior tip of the upper jaw (premaxilla) to the posterior end of the last vertebra (hypural plate).

### RNA extraction and sequencing

We removed frozen tissues from -80℃ and allowed them to thaw gradually by first putting them in a refrigerator for approximately 30 minutes before exposing them to room temperature. For each tissue, we cut about 3.5-4.5mg for extraction and the rest of the tissue was stored in a -80℃ freezer. We used the Qiagen RNAeasy kit (RNeasy Mini Kit (50), Cat no.74104) to extract the RNA from tissues by following the manufacturer instructions with slight modifications. We added 2-Mercaptoethanol (1ML 2-Mercaptoethanol, Cat No: 97622, Sigma Aldrich) to the lysis buffer. After extraction, we quantified the RNA concentration in each sample using the Qubit Fluorometer 3.0 set to the High Sensitivity option for RNA. The concentrations of the RNA were normalized within the range required by the Genomic Sequencing and Analysis Facility of University of Texas at Austin. The minimum concentration of the 30 specimens was 1.7 ng/ul while the maximum was 82.7ng/ul. The samples were sent to the Genomic Sequencing and Analysis Facility of University of Texas at Austin for library preparation and RNA sequencing on the Illumina NovaSeq 6000 SR100.

### TagSeq

We sequenced the extracted RNA using a revised version of the TagSeq approach(Lohman et al. 2016). TagSeq involves sequencing only the 3’ end of mRNAs, which reduces the cost compared to whole mRNA sequencing. This approach increases the power of an experimental design as more samples can be sequenced at relatively low cost. However, this increased power comes at the expense of reduced information. In particular, TagSeq is unable to distinguish alternatively spliced variants from one genomic locus, as well as polymorphisms within the gene locus. Nevertheless, this approach fits the goal of this study, which is to leverage existing genetic variation information for various populations of threespine stickleback, including from Rabbit Slough and Cheney Lake, two of the populations used in this study.

### Bioinformatic processing

We adapted the pipeline for processing TagSeq raw reads from https://github.com/z0on/tag-based_RNAseq. We concatenated read files belonging to the same specimen and tissue (each specimen had brain and gill tissues) and trimmed adapters using the software cutadapt v1.14(Martin 2011) with Python 2.7.15 and command line parameters: - -a AAAAAAAA -a AGATCGG -q 15 -m 25. We then aligned the reads to the stickleback reference genome stickleback_v5 (https://stickleback.genetics.uga.edu/downloadData/v5_assembly/stickleback_v5_assembly.fa.gz)(Peichel et al. 2020) using bowtie2 with the --local option. We used the script samcount.pl (https://github.com/z0on/tag-based_RNAseq/blob/master/samcount.pl) to count isoforms mapping to each gene.

### Processing of bulk RNAseq

In order to compare our newly generated TagSeq data to existing, comparable bulk RNAseq, we reprocessed a subset of the Verta and Jones 2019(Verta & Jones 2019) dataset from the gill, including three marine and three freshwater individuals collected from Little Campbell River, British Columbia, Canada. The marine samples were collected from 49.016, -122.780, while the freshwater individuals were collected from 49.012, -122.625. There were one male and two female marine individuals, and two females and one male freshwater individuals. We downloaded the reads from SRA (accession numbers: SRR8868247, SRR8868248, SRR8868249, SRR8868250, SRR8868251, SRR8868252, SRR8868254). Subsequently, we performed adapter removal using trim-galore. We then mapped the reads to the stickleback reference genome stickleback_v5 (https://stickleback.genetics.uga.edu/downloadData/v5_assembly/stickleback_v5_assembly.fa.gz)(Peichel et al. 2020) and counted the number of reads per gene using STAR(Dobin et al. 2013). Similar to Verta and Jones 2019(Verta & Jones 2019), we used the following settings in STAR aligner: --readFilesCommand zcat --outSAMtype BAM SortedByCoordinate --quantMode GeneCounts --outFilterIntronMotifs RemoveNoncanonicalUnannotated --chimSegmentMin 50 --alignSJDBoverhangMin 1 --alignIntronMin 20 --alignIntronMax 200000 –alignMatesGapMax 200000 --limitSjdbInsertNsj 2000000.

### Differential gene expression

All samples had more than one million reads, with the exception of one brain sample from Rabbit Slough. We therefore removed this sample from all downstream analyses. We used DESeq2(Love et al. 2014) to perform normalization on the count data before performing differential gene expression. DESeq2 models count data using a negative binomial distribution and apply median-of-ratios normalization(Anders & Huber 2010). To reduce noise from lowly expressed transcripts and improve statistical power in downstream analyses, we removed all genes with base mean count of less than 10 across all samples. We performed differential gene expression analysis between marine and freshwater populations separately for the brain and gill tissues.

### Genomic datasets and processing

In order to contextualize RNA expression with patterns of genetic variation, we utilized genomic data from a 2009 Rabbit Slough sample previously published by Roberts Kingman et al. 2021(Roberts Kingman et al. 2021), while genomes from Cheney were collected in June 2020 and were previously published by (Kwakye et al. 2025). For Cornelius Lake, no previous genomic dataset existed, therefore we generated genomic data using the five individuals we sampled for the RNA-seq. Bioinformatic processing of the Cornelius Lake genomes followed the same pipeline previously used in (Roberts Kingman et al. 2021) and (Kwakye et al. 2025). Briefly, we extracted DNA samples using the Qiagen DNeasy 96 Blood & Tissue Kit for animal tissue and quantified them with a Qubit Fluorometer 3.0 set to the High Sensitivity option for dsDNA. We performed genome resequencing at Beijing Genomics Institute (BG) using their proprietary DNBseq technology. We trimmed adapters with AdapterRemoval (ver. 2.2.2)(Lindgreen 2012), and then mapped reads to the Threespine Stickleback genome version gasAcu1-4(Peichel et al. 2017; Roberts Kingman et al. 2021) using bwa mem -M (v0.7.15-r1140)(Li & Durbin 2009). We added read groups using Picard before being merged with samtools(Danecek et al. 2021). We also marked duplicates using Picardtools. We performed base recalibration using BaseRecalibrator from GATK ver3.7(McKenna et al. 2010; Van Der Auwera et al. 2013).

### Population structure and signatures of selection

Due to the relatively lower mean coverage of the Cornelius Lake individuals (mean coverage of approximately 6X compared to the approximately 20X coverage for Rabbit Slough and 30X for Cheney Lake), we performed pairwise *F_ST_* using angsd(Korneliussen et al. 2014), which is designed for low coverage genomic data. We first estimated the site allele frequency likelihood for each population using angsd (using 20 individuals from Rabbit Slough, 20 individuals from Cheney Lake and 5 individuals from Cornelius Lake). We then calculated the 2d site frequency spectrum (SFS) between each population pair (Rabbit Slough vs Cheney Lake, Rabbit Slough vs Cornelius Lake and Cheney Lake vs Cornelius Lake) using realSFS. The pairwise F_ST_ was then estimated using the site allele frequency likelihood and the 2d sfs. We also performed principal component analyses (PCA) using a combination of angsd and pcangsd(Meisner & Albrechtsen 2018). Again, we used angsd to estimate the genotype likelihoods, but using all individuals in the three populations combined (n=45). We then used pcangsd to estimate the covariance matrix and the eigen function in *R* to compute the eigenvalues and eigenvectors.

### Enrichment of signatures of selection in differentially expressed genes (DEGs)

We used bedtools intersect to find the SNPs located in DEGs. To capture putative *cis*-regulatory regions, including promoters and enhancers, we included SNPs found within 10Kbp up and downstream of the gene. The distance between enhancers of genes and the promoters they interact with in eukaryotic organisms can vary from a few Kbp to 1Mbp, with certain enhancers found in other genes(Ritter et al. 2012; Birnbaum et al. 2012). While a 10Kbp threshold does not capture long range enhancers, this threshold serves as a balance between gene specific SNPs and those that can overlap multiple other genes. We determined the enrichment of F_ST_ outliers (we defined F_ST_ outliers as values greater than 99^th^ percentile, see below) in DEGs compared to the rest of the genome using Fisher’s Exact Test.

### Genetic load

We annotated the multi-populationVCF using snpEff (version SnpEff 4.3t (build 2017-11-24 10:18))(Cingolani et al. 2012). We built a custom database for the stickleback genome gascu1-4(Roberts Kingman et al. 2021) using the following command: java -jar snpEff.jar build -gtf22 - v Gascu_1-4 -noCheckCds -noCheckProtein. We then ran snpEff annotation with and without the -noShiftHgvs option. As we mapped the RNA-seq data to the v5 of stickleback genome, we lifted over the bed files of all differentially expressed genes from the v5 to v4 using liftOver and chain files from https://stickleback.genetics.uga.edu/downloadData/chain_files/v5_to_v4.chain.txt. We also determined genetic load across the DEGs using GERP scores. For this analysis, we downloaded conservation scores (https://ftp.ensembl.org/pub/release-112/compara/conservation_scores/65_fish.gerp_conservation_score/) for the 65 fish multiple alignment (https://ftp.ensembl.org/pub/release-112/emf/ensembl-compara/multiple_alignments/65_fish.epo_extended/) from the Ensembl release 112. We then lifted over these sites from version 5 (https://ftp.ensembl.org/pub/release-112/fasta/gasterosteus_aculeatus/dna/Gasterosteus_aculeatus.GAculeatus_UGA_version5.dna_rm.toplevel.fa.gz) to version gascu1-4 using liftOver and chain files from https://stickleback.genetics.uga.edu/downloadData/chain_files/v5_to_v4.chain.txt.

## Author Contributions

A.K: experimental design, sampling, formal analysis, RNA and DNA extraction and quantification, library preparation, data processing; N.D: RNA and DNA extraction and quantification, M.W: sampling; K.R.V: experimental design, project supervision. A.K wrote the initial draft with input from K.R.V and M.W. All authors contributed to production of the final version.

## Acknowledgments

This work was supported by NIH R01GM124330 to K.R.V. and institutional funding from The College of New Jersey to M.A.W.). We appreciate Alison Bell for comments on an early version of this manuscript. We thank members of Wund Lab (Neel Patel and Sathya Rameshkumar) for their assistance during the field trip. We also thank members of the Veeramah Lab for their comments and discussions during the preparation of this manuscript.

