## Supplementary Figures and Tables for "Genomic signatures of selection are enriched in differentially expressed genes in sticklebacks adapting to contrasting environments"

**Fig. S1** Genotype at regions of the genome identified by (Roberts Kingman et al. 2021) to be underlying rapid freshwater adaptation for **A)** Rabbit Slough **B)** Cheney Lake **C)** Cornelius Lake samples.

**Fig. S2**: Proportion of freshwater adaptive alleles for individuals collected from Rabbit Slough, Cheney Lake, Cornelius Lake, and genomes from freshwater populations published in Roberts Kingmann et al. (Roberts Kingman et al. 2021)

**Fig. S3** Correlation of gene counts estimated using TagSeq for genes expressed in the brain for Rabbit Slough individuals vs genes expressed in the gill for marine individuals collected from Little Campbell River

**Fig. S4: Correlation of gene expression between marine and freshwater stickleback populations.** Correlation analysis of normalized transcript counts in gill tissue between Cheney and Rabbit Slough populations. Each point represents an individual gene. Differentially expressed genes (DEGs) are highlighted in red and housekeeping genes in blue.

**Fig. S5 Correlation of gene expression between marine and freshwater stickleback populations.** Correlation analysis of normalized transcript counts in gill tissue between Cornelius and Rabbit Slough populations. Each point represents an individual gene. Differentially expressed genes (DEGs) are highlighted in red and housekeeping genes in blue.

**Fig. S6 Correlation of gene expression between marine and freshwater stickleback populations.** Correlation analysis of normalized transcript counts in gill tissue between marine and freshwater populations from Verta and Jones 2019. Each point represents an individual gene. Differentially expressed genes (DEGs) are highlighted in red and housekeeping genes in blue.

**Fig. S7** Genome-wide distribution of GERP scores

**Fig. S8** Distribution of GERP scores for *dhx58*

**Fig. S9** Distribution of GERP scores for *pvalb4*

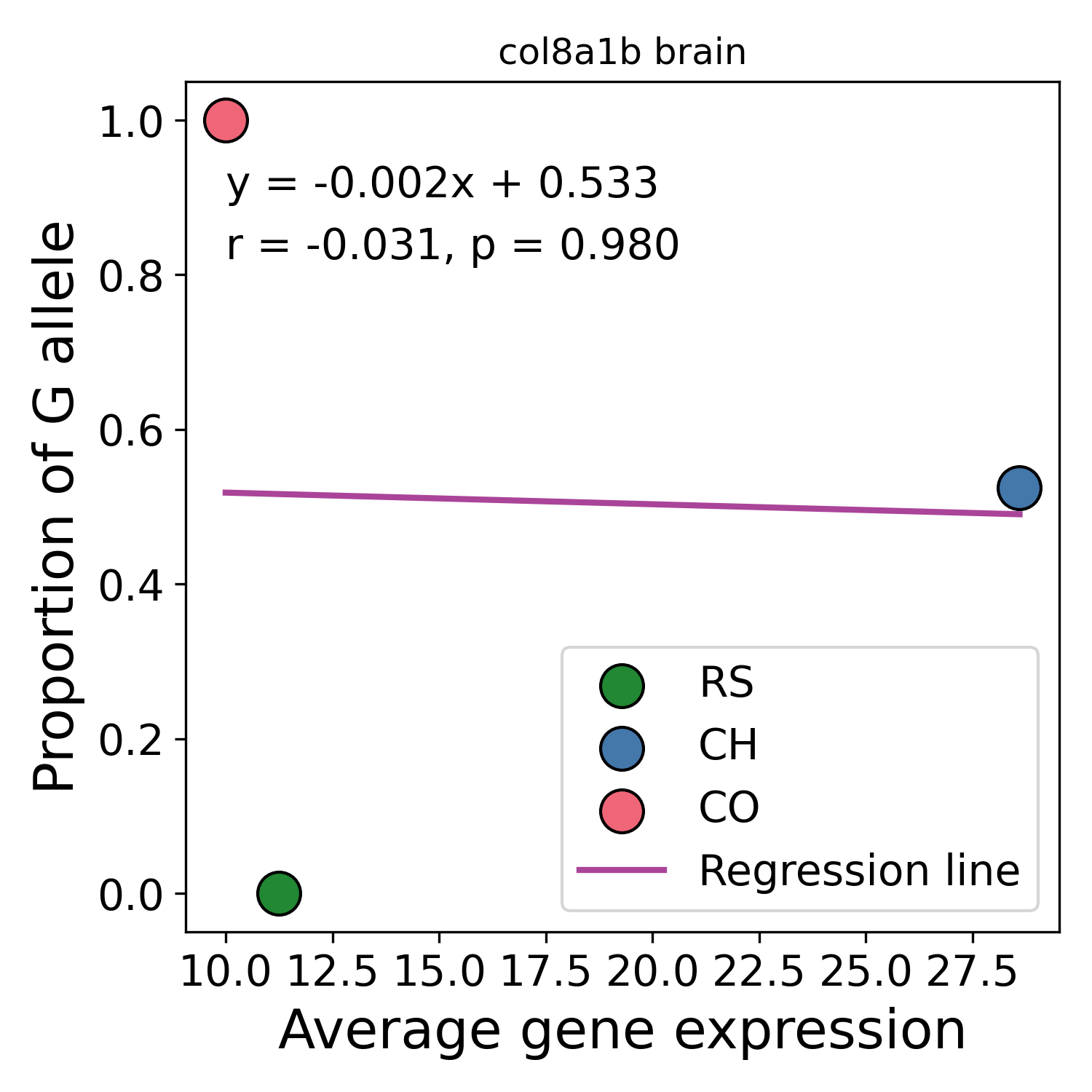

Fig. S10: Correlation between the average gene expression in the brain tissue and the proportion of G alleles at a SNP((chrI:26106268 G > T) predicted to have a high impact on the gene *col8a1b*.

**Supplementary Tables**

*Table S1: Gill DEGs found in Rabbit Slough vs Cheney Lake F_ST_ outliers*

| Gene name | Gene ID | Chromosome | FST outliers | Tempo SNPs |
| --- | --- | --- | --- | --- |
| *fosb* | ENSGACG00000009792 | chrI | 2 |  |
| *cahz* | ENSGACG00000014774 | chrIII | 22 |  |
| *col6a1* | ENSGACG00000012576 | chrIII | 18 |  |
| *cmah* | ENSGACG00000019485 | chrIV | 49 |  |
| *tnmd* | ENSGACG00000017466 | chrIV | 3 |  |
| *hbegfa* | ENSGACG00000018355 | chrIV | 29 |  |
| *antxr1c* | ENSGACG00000004906 | chrV | 1 |  |
| *ENSGACG00000002171* | ENSGACG00000002171 | chrVIII | 1 |  |
| *ptgs2b* | ENSGACG00000007384 | chrVIII | 21 |  |
| *col11a2* | ENSGACG00000000099 | chrX | 1 |  |
| ***pvalb4*** | **ENSGACG00000012174** | **chrXI** | **124** | **30** |
| *tnnt1* | ENSGACG00000006626 | chrXI | 7 |  |
| *mylpfa* | ENSGACG00000013640 | chrXI | 1 |  |
| *cacnb1* | ENSGACG00000009291 | chrXI | 139 |  |
| *itga5* | ENSGACG00000010945 | chrXII | 30 |  |
| *si:dkey-205h13.1* | ENSGACG00000011044 | chrXII | 20 |  |
| *bgnb* | ENSGACG00000010078 | chrXII | 10 |  |
| *si:dkey-245n4.2* | ENSGACG00000010369 | chrXIII | 10 |  |
| *ENSGACG00000006275* | ENSGACG00000006275 | chrXVI | 1 |  |
| *ldha* | ENSGACG00000011270 | chrXIX | 3 |  |
| *rbm24b* | ENSGACG00000008472 | chrXX | 6 |  |
| *hsc70* | ENSGACG00000007569 | chrXX | 12 |  |
| *gapdh* | ENSGACG00000010219 | chrXX | 17 |  |
| *pi15a* | ENSGACG00000002723 | chrXXI | 19 |  |

*Table S2: Gill DEGs found in Rabbit Slough vs Cornelius Lake F_ST_ outliers*

| Gene name | Gene ID | Chromosome | F_ST_ outliers | Tempo SNPs |
| --- | --- | --- | --- | --- |
| ***col8a1b*** | **ENSGACG00000014373** | **chrI** | **227** | **40** |
| *col6a1* | ENSGACG00000012576 | chrIII | 7 |  |
| *nxpe3* | ENSGACG00000019508 | chrIV | 2 |  |
| *cmah* | ENSGACG00000019485 | chrIV | 30 |  |
| ***tmem168a*** | **ENSGACG00000018918** | **chrIV** | **57** | **18** |
| *pcolcea* | ENSGACG00000020298 | chrVII | 79 |  |
| *S100P* | ENSGACG00000020061 | chrVII | 1 |  |
| ***si:ch211-156l18.7*** | **ENSGACG00000018299** | **chrIX** | **79** | **13** |
| ***sod3b*** | **ENSGACG00000017840** | **chrIX** | **93** | **26** |
| *myoz2b* | ENSGACG00000017804 | chrIX | 116 |  |
| *slc25a4* | ENSGACG00000017708 | chrIX | 85 |  |
| *ST3GAL1* | ENSGACG00000008844 | chrX | 1 |  |
| ***pvalb4*** | **ENSGACG00000012174** | **chrXI** | **94** | **28** |
| *ENSGACG00000005444* | ENSGACG00000005444 | chrXI | 1 |  |
| ***pvalb8*** | **ENSGACG00000012178** | **chrXI** | **88** | **27** |
| *ENSGACG00000012669* | ENSGACG00000012669 | chrXI | 2 |  |
| *ENSGACG00000012690* | ENSGACG00000012690 | chrXI | 4 |  |
| ***si:dkey-205h13.1*** | **ENSGACG00000011044** | **chrXII** | **284** | **59** |
| *itga5* | ENSGACG00000010945 | chrXII | 1 |  |
| *bgnb* | ENSGACG00000010078 | chrXII | 1 |  |
| ***si:dkey-245n4.2*** | **ENSGACG00000010369** | **chrXIII** | **124** | **8** |
| ***nol6*** | **ENSGACG00000010296** | **chrXIII** | **312** | **8** |
| *ldha* | ENSGACG00000011270 | chrXIX | 7 |  |
| *atp6v1e1a* | ENSGACG00000013695 | chrXIX | 3 |  |
| *calb2a* | ENSGACG00000003013 | chrXIX | 2 |  |
| *celf3b* | ENSGACG00000011707 | chrXX | 185 |  |
| *tubb5* | ENSGACG00000009635 | chrXX | 5 |  |
| ***hsc70*** | **ENSGACG00000007569** | **chrXX** | **184** | **35** |
| ***hgd*** | **ENSGACG00000002559** | **chrXXI** | **429** | **8** |

*Table S3:Brain DEGs found in only Rabbit Slough vs Cheney Lake F_ST_ outliers*

| Gene name | Gene ID | Chromosome | FST outliers | Tempo SNPs |
| --- | --- | --- | --- | --- |
| *MAP4K1* | ENSGACG00000013172 | chrI | 4 |  |
| *cahz* | ENSGACG00000014774 | chrIII | 22 |  |
| *col6a1* | ENSGACG00000012576 | chrIII | 18 |  |
| *slc7a11* | ENSGACG00000019609 | chrIV | 14 |  |
| *CPNE8* | ENSGACG00000019027 | chrIV | 28 |  |
| *strip2* | ENSGACG00000018938 | chrIV | 13 |  |
| *foxp2* | ENSGACG00000018911 | chrIV | 4 |  |
| *ENSGACG00000018550* | ENSGACG00000018550 | chrIV | 4 |  |
| ***ENSGACG00000018361*** | **ENSGACG00000018361** | **chrIV** | **161** | **27** |
| ***n4bp3*** | **ENSGACG00000018359** | **chrIV** | **105** | **40** |
| *hbegfa* | ENSGACG00000018355 | chrIV | 29 |  |
| *ENSGACG00000018050* | ENSGACG00000018050 | chrIV | 3 |  |
| ***DOCK2*** | **ENSGACG00000017885** | **chrIV** | **216** | **39** |
| *ponzr1* | ENSGACG00000019324 | chrVII | 4 |  |
| *pnp5a* | ENSGACG00000018988 | chrVII | 2 |  |
| *myo18aa* | ENSGACG00000020305 | chrVII | 7 |  |
| *ENSGACG00000020069* | ENSGACG00000020069 | chrVII | 19 |  |
| *S100P* | ENSGACG00000020061 | chrVII | 8 |  |
| *ENSGACG00000018560* | ENSGACG00000018560 | chrVII | 2 |  |
| *f11r.1* | ENSGACG00000020422 | chrVII | 17 |  |
| *ENSGACG00000007974* | ENSGACG00000007974 | chrVIII | 5 |  |
| *slc25a24* | ENSGACG00000006388 | chrVIII | 2 |  |
| *b3gnt3.4* | ENSGACG00000008643 | chrVIII | 11 |  |
| *ENSGACG00000002171* | ENSGACG00000002171 | chrVIII | 1 |  |
| *add3a* | ENSGACG00000018717 | chrIX | 3 |  |
| *capn2b* | ENSGACG00000019944 | chrIX | 2 |  |
| *ENSGACG00000018430* | ENSGACG00000018430 | chrIX | 1 |  |
| *ENSGACG00000006358* | ENSGACG00000006358 | chrX | 1 |  |
| *ENSGACG00000014556* | ENSGACG00000014556 | chrXI | 1 |  |
| *slc4a1a* | ENSGACG00000009622 | chrXI | 8 |  |
| ***dhx58*** | **ENSGACG00000008740** | **chrXI** | **153** | **80** |
| *PKP1* | ENSGACG00000009752 | chrXII | 19 |  |
| *galnt6* | ENSGACG00000008590 | chrXII | 1 |  |
| *si:dkey-205h13.1* | ENSGACG00000011044 | chrXII | 20 |  |
| *ENSGACG00000011026* | ENSGACG00000011026 | chrXIII | 4 |  |
| *ENSGACG00000006275* | ENSGACG00000006275 | chrXVI | 1 |  |
| *rhag* | ENSGACG00000009865 | chrXVIII | 1 |  |
| *ENSGACG00000011188* | ENSGACG00000011188 | chrXIX | 4 |  |
| *nbeal2* | ENSGACG00000008637 | chrXX | 14 |  |
| *ENSGACG00000006045* | ENSGACG00000006045 | chrXX | 26 |  |
| *ncam3* | ENSGACG00000005994 | chrXX | 7 |  |
| *ENSGACG00000012665* | ENSGACG00000012665 | chrXX | 1 |  |
| *stmn1b* | ENSGACG00000007379 | chrXX | 9 |  |
| *tmod4* | ENSGACG00000013516 | chrXX | 1 |  |
| *arfgef1* | ENSGACG00000002879 | chrXXI | 85 |  |
| *tram1* | ENSGACG00000002818 | chrXXI | 46 |  |
| *cmbl* | ENSGACG00000002636 | chrXXI | 37 |  |

*Table S4:Brain DEGs found in only Rabbit Slough vs Cornelius Lake F_ST_ outliers*

| Gene name | Gene ID | Chromosome | F_ST_ outliers | Tempo SNPs |
| --- | --- | --- | --- | --- |
| *arrb1* | ENSGACG00000004570 | chrI | 21 |  |
| ***ENSGACG00000018030*** | **ENSGACG00000018030** | **chrIV** | **125** | **18** |
| *slc6a1l* | ENSGACG00000019748 | chrIV | 29 |  |
| *slc7a11* | ENSGACG00000019609 | chrIV | 1 |  |
| *pfkfb3* | ENSGACG00000019176 | chrIV | 1 |  |
| *SYN3* | ENSGACG00000019125 | chrIV | 350 |  |
| ***strip2*** | **ENSGACG00000018938** | **chrIV** | **100** | **5** |
| ***ITM2A*** | **ENSGACG00000018559** | **chrIV** | **131** | **55** |
| ***acsl4a*** | **ENSGACG00000018503** | **chrIV** | **274** | **82** |
| *ENSGACG00000018050* | ENSGACG00000018050 | chrIV | 8 |  |
| *vamp2* | ENSGACG00000020297 | chrVII | 62 |  |
| ***ENSGACG00000020330*** | **ENSGACG00000020330** | **chrVII** | **125** | **79** |
| ***ENSGACG00000020069*** | **ENSGACG00000020069** | **chrVII** | **217** | **102** |
| *cacng1b* | ENSGACG00000012215 | chrXI | 1 |  |
| ***dhx58*** | **ENSGACG00000008740** | **chrXI** | **50** | **31** |
| ***si:dkey-205h13.1*** | **ENSGACG00000011044** | **chrXII** | **284** | **59** |
| *csad* | ENSGACG00000009463 | chrXII | 12 |  |
| ***si:dkey-245n4.2*** | **ENSGACG00000010369** | **chrXIII** | **124** | **8** |
| *ENSGACG00000008429* | ENSGACG00000008429 | chrXIII | 23 |  |
| *ENSGACG00000012351* | ENSGACG00000012351 | chrXX | 3 |  |
| ***arfgef1*** | **ENSGACG00000002879** | **chrXXI** | **795** | **13** |
| ***gdap1*** | **ENSGACG00000002727** | **chrXXI** | **303** | **2** |
| ***cmbl*** | **ENSGACG00000002636** | **chrXXI** | **342** | **6** |

*Table S5: Missense variants associated with DEGs*

| Chromosome | Position | Ref | Alt | Gene |
| --- | --- | --- | --- | --- |
| **Rabbit Slough vs Cheney Brain DEGs** | | | | |
| chrIV | 13353322 | A | G | *n4bp3* |
| chrIV | 13373668 | T | A |  |
| chrIV | 13373744 | T | C |  |
| chrXI | 6629831 | A | T | *kat2a* |
| chrXI | 6636856 | A | C | *dhx58* |
| chrXI | 6637178 | G | A | *dhx58* |
| chrXI | 6639650 | A | G | *dhx58* |
| **Rabbit Slough vs Cornelius Brain DEGs** | | | | |
| chrIV | 11016657 | T | G | *myoz3a* |
| chrIV | 15071531 | C | G | *acsl4a* |
| chrIV | 15073110 | C | A | *acsl4a* |
| chrIV | 15073356 | A | G | *acsl4a* |
| chrIV | 15073383 | A | G | *acsl4a* |
| chrIV | 15076140 | T | A | *acsl4a* |
| chrIV | 15745378 | T | G | *gpr174* |
| chrIV | 15745501 | T | C | *gpr174* |
| chrIV | 15745531 | A | C | *gpr174* |
| chrIV | 15745773 | T | G | *gpr174* |
| chrIV | 15745775 | T | C | *gpr174* |
| chrIV | 26923560 | T | C | *strip2* |
| chrVII | 10849566 | G | A | *tbc1d14* |
| chrVII | 10858813 | C | A |  |
| chrXI | 6636856 | A | C | *dhx58* |
| chrXII | 6616188 | G | A | *si:dkey-205h13.1* |
| chrXIII | 10698847 | T | G | *aqp3a* |
| **Rabbit Slough vs Cornelius Gill DEGs** | | | | |
| chrI | 26106349 | C | G | *col8a1b* |
| chrI | 26106391 | T | G | *col8a1b* |
| chrI | 26106589 | G | T | *col8a1b* |
| chrI | 26106634 | G | C | *col8a1b* |
| chrI | 26106679 | T | C | *col8a1b* |
| chrI | 26106727 | T | A | *col8a1b* |
| chrI | 26106956 | C | A | *col8a1b* |
| chrI | 26107096 | T | C | *col8a1b* |
| chrI | 26107789 | G | C | *col8a1b* |
| chrI | 26108583 | A | T | *col8a1b* |
| chrI | 26108590 | T | C | *col8a1b* |
| chrI | 26108617 | T | C | *col8a1b* |
| chrI | 26108692 | C | T | *col8a1b* |
| chrIX | 11316900 | G | C | *si:ch211-156l18.7* |
| chrIX | 11318337 | C | T | *si:ch211-156l18.7* |
| chrIX | 13523523 | C | G | *sod3b* |
| chrIX | 13523602 | A | G | *sod3b* |
| chrIX | 13523614 | A | G | *sod3b* |
| chrIX | 13527072 | C | T | *DHX15_(1_of_many)* |
| chrIX | 13528335 | T | C | *DHX15_(1_of_many)* |
| chrXII | 6616188 | G | A | *si:dkey-205h13.1* |
| chrXIII | 10698847 | T | G | *aqp3a* |
| chrXX | 10614960 | G | A | *lin37* |
| chrXX | 10618297 | G | C | *si:ch211-137j23.7* |
| chrXX | 10618381 | A | G | *si:ch211-137j23.7* |
| chrXX | 10618408 | A | G | *si:ch211-137j23.7* |
| chrXX | 10618602 | T | C | *si:ch211-137j23.7* |
| chrXX | 10618636 | C | A | *si:ch211-137j23.7* |
| chrXX | 10618651 | G | A | *si:ch211-137j23.7* |
| chrXX | 10623887 | G | T | *zgc:171592* |
